# IGF-1 facilitates extinction of conditioned fear

**DOI:** 10.1101/2020.04.15.042788

**Authors:** José A. Noriega-Prieto, Laura E. Maglio, Irene B. Maroto, Jesús Martin-Cortecero, Antonio Muñoz-Callejas, Marta Callejo-Móstoles, David Fernández de Sevilla

**Affiliations:** Departamento de Anatomía, Histología y Neurociencia, Facultad de Medicina, Universidad Autónoma de Madrid, 28029 Madrid, Spain; Department of Neuroscience, University of Minnesota, Minneapolis, MN 55455, USA; Departamento de Ciencias Médicas Básicas (Fisiología) and Instituto de Tecnologías Biomédicas (ITB), Universidad de La Laguna, 38071 Tenerife, Spain; Centro de Investigación Biomédica en Red Sobre Enfermedades Neurodegenerativas (CIBERNED), Instituto Universitario de Investigación Neuroquímica (IUIN), Instituto Ramón y Cajal de Investigación Sanitaria (IRYCIS) and Departmento de Bioquímica y Biología Molecular, Facultad de Química, Universidad Complutense de Madrid, 28040 Madrid, Spain; Institute of Physiology and Pathophysiology, Heidelberg University, 69120 Heidelberg, Germany

**Author notes:** These authors contributed equally (José A. Noriega-Prieto, Laura E. Maglio). Corresponding authors, Correspondence should be addressed to: David Fernández de Sevilla, PhD, Departamento de Anatomía, Histología y Neurociencia, Facultad de Medicina, Universidad Autónoma de Madrid, c/ Arzobispo Morcillo, 4, 28029, Madrid, Spain., Laura Eva Maglio, PhD, Departamento de Ciencias Médicas, Básicas (Fisiología), Facultad de Medicina, Universidad de La Laguna, Campus de Ofra s/n, 38071 San, Cristóbal de La Laguna, Tenerife, Spain.

**Keywords:** IGF-1, Extinction of condition fear, afterhyperpolarization, synaptic plasticity, infralimbic cortex

## Abstract

Insulin-like growth factor-1 (IGF-1) plays a key role in synaptic plasticity, degenerative diseases, spatial learning, and anxiety-like behavioral processes. While IGF-1 regulates neuronal activity in many areas of the brain, its effect on synaptic plasticity and animal behavior dependent on the prefrontal cortex remain unexplored. Here, we show that IGF-1 induces a long-lasting depression of the medium and slow post-spike afterhyperpolarization (mAHP and sAHP), increasing the excitability of layer 5 pyramidal neurons of the infralimbic cortex. Besides, IGF-1 mediates a long-term depression of both inhibitory and excitatory synaptic transmission that results in a longterm potentiation of the postsynaptic potentials. We demonstrate that these synaptic and intrinsic regulatory processes mediated by IGF-1 favor the fear extinction memory. These results show novel functional consequences of IGF-1 signaling on animal behavior tasks dependent on the prefrontal cortex, revealing IGF-1 as a key element in the control of the fear extinction memory.

**Impact Statement:** IGF-1 modulates the neuronal firing and synaptic plasticity in infralimbic cortex, favoring the extinction memory

## Introduction

Conditioned fear is an associative form of learning and memory in which a previous neutral stimulus (called “conditioned stimulus” [CS]; e.g., a tone) comes to elicit a fear response when is associated with an aversive stimulus (called “unconditioned stimulus” [US]; e.g., an electric shock (Ledoux, 2000; Maren, 2001; Pape and Pare, 2010). Extinction of conditioned fear is an active learning process involving inhibition of fear expression. It is a decline in conditioned fear responses (CRs) following non-reinforced exposure to the CS. However, fear extinction memory does not erase the initial association between the CS-US but forms a new association (CS-No US) that inhibits expression of the previous conditioned memory (Quirk and Mueller, 2008). Fear extinction depends on specific structures such as the amygdala, the hippocampus, and the prefrontal cortex (PFC) (Milad and Quirk, 2012; Orsini and Maren, 2012). Dysfunctions in the neuronal circuits responsible for fear cause the development of anxiety disorders, including specific phobias and post-traumatic stress (Pavlov, 1927; Quirk and Mueller, 2008).

The consolidation of the extinction memory has been related to long-term synaptic plasticity and increases in the excitability of pyramidal neurons (PNs) from the infralimbic cortex (IL) (Kaczorowski et al., 2012; Koppensteiner et al., 2019; Moyer et al., 1996). The mechanisms of this synaptic plasticity involve NMDA receptors (Burgos-Robles et al., 2007), mitogen-activated protein (MAP) kinases (Hugues et al., 2004), protein kinase A (Mueller et al., 2008), insertion of Ca^2+^-permeable AMPA receptors (Sepulveda-Orengo et al., 2013), and protein synthesis (Mueller et al., 2008; Santini et al., 2004). Moreover, the excitability of PN from IL is a key determinant for both the acquisition and the extinction of fear, being reduced by fear acquisition and increased by fear extinction (Santini et al., 2008). Indeed, IL layer 5 pyramidal neurons (L5PN) of conditioned fear animals show a higher slow post-spike afterhyperpolarization (sAHP) amplitude and lower firing frequency relative to non-conditioned or extinguished animals (Santini et al., 2008).

Insulin-like growth factor-1 (IGF-1) is an endogenous polypeptide with plenty of trophic functions, which can be locally synthesized and released by neurons and astrocytes (Fernandez and Torres-Alemán, 2012). Similarly, its receptor (IGF-1R) is widely expressed among all brain cell type (Quesada, et al., 2007; Rodriguez-Perez et al., 2016). IGF-1 increases neuronal firing (Gazit et al., 2016; Nuñez et al., 2003) and modulates excitatory and inhibitory synaptic transmission in the central nervous system (Castro-alamancos and Torres-aleman, 1993; Nilsson et al., 1988; Noriega-prieto et al., 2020; Seto et al., 2002). Furthermore, IGF-1 regulates different ion channels, such as A-type K^+^ channels (Xing et al., 2006) and P/Q-, L-, and N-type voltage-gated Ca^2+^ channels (Blair and Marshall, 1997; Shan et al., 2003), as well as neurotransmitter receptors activity (Gonzalez de la Vega et al., 2001; Savchenko et al., 2001). However, the role of IGF-1 on the modulation of L5PN of the IL activity and its consequences in the fear extinction memory remains to be clarified.

Here, we have examined the effects of IGF-1 in excitability and synaptic transmission in L5PN from IL. Our results reveal that IGF-1 induces a long-lasting reduction of the sAHP amplitude and increases the firing frequency of L5PNs. Furthermore, IGF-1 induces a presynaptic long-term depression (LTD) of both excitatory and inhibitory postsynaptic currents (EPSC and IPSCs, respectively) that results in a long-term potentiation (LTP) of the postsynaptic potentials (PSPs). Moreover, we show that IGF-1 facilitates the recall of extinction of fear conditioning when applied intracranially 30 min before the extinction task. In these animals with the favored extinction, the sAHP is reduced and the excitatory synaptic transmission is depressed. Therefore, we demonstrate for the first time that IGF-1 facilitates the extinction of fear conditioning, increasing the excitability and potentiating the synaptic transmission in L5PN from IL.

## Materials And Methods

### Animals

Male Sprague Dawley rats were group-housed in transparent polyethylene cages. Rats were maintained on a 12:12 h light/dark scheduled cycle with free access to food and water. All animal procedures were approved by the Universidad Autónoma of Madrid Ethical Committee on Animal Welfare and conform to Spanish and European guidelines for the protection of experimental animals (Directive 2010/63/EU). Effort was made to minimize animal suffering and number.

### Electrophysiology

Prefrontal cortical slices were obtained from rats at postnatal day (P20 to P30) age. Rats were decapitated and the brain removed and submerged in artificial cerebrospinal fluid (ACSF). Coronal slices (400 μm thick) were obtained with a Vibratome (Leica VT 1200S). To reduce swelling and damage in superficial layers (especially after the behavior tests), brain slices were obtained using a modified ACSF, containing (in mM): 75 NaCl, 2.69 KCl, 1.25 KH2PO4, 2 MgSO4, 26 NaHCO3, 2 CaCl2, 10 glucose, 100 sucrose, 1 sodium ascorbate, and 3 sodium pyruvate. After that, brain slices were transferred to regular ACSF (in mM: 124 NaCl, 2.69 KCl, 1.25 KH_2_PO_4_, 2 Mg _2_SO_4_, 26 NaHCO_3_, 2 CaCl_2_, and 10 glucose, 0.4 sodium ascorbate, bubbled with carbogen [95% O_2_, 5% CO_2_]) and incubated for >1h at room temperature (22-24°C). Slices were then transferred to an immersion recording chamber and superfused with carbogen-bubbled ACSF. Cells were visualized under an Olympus BX50WI microscope. Patch-clamp recordings were obtained from the soma of pyramidal neurons located at the layer 5 of IL (IL-L5PNs) using patch pipettes (4-8MΩ) filled with an internal solution that contained (in mM): 130 KMeSO_4_, 10 HEPES-K, 4 Na_2_ATP, 0.3 Na_3_GTP, 0.2 EGTA, 10 KCl (buffered to pH 7.2-7.3 with KOH). Recordings were performed in current-or voltage-clamp modes using a Cornerstone PC-ONE amplifier (Dagan Corporation). Pipettes were placed with a micromanipulator (Narishige). The holding potential was adjusted to −60 mV and the series resistance was compensated to ~80%. Recordings were accepted only when series and input resistances did not change >20% during the experiment. Data were low-pass filtered at 3 kHz and sampled at 10 kHz, through a Digidata 1440 (Molecular Devices). The pClamp software (Molecular Devices) was used to acquire the data. Chemicals were purchased from Sigma-Aldrich Quimica and Tocris Bioscience (Ellisville; distributed by Biogen Cientifica) and R&D Systems, Inc. (distributed by Bio-Techne). In current-clamp mode, we examined the afterhyperpolarizing potentials (AHPs) in IL-L5PNs. We injected 0.4 nA depolarizing current pulses of 10, 200, and 800 ms in duration to record fAHPs, mAHPs, and sAHPs, respectively. We measured the amplitude of afterhyperpolarizing potentials as the average of the maximum peak value, after the end of each depolarized pulse. The temporal window of each one was 5-10 ms for fAHPs, 5-100 ms for mAHPs, and 0.1-3 s for sAHPs, measured in ACSF and in the presence of 10 nM IGF-1. We applied the same protocol but in the voltage-clamp mode to record the currents that underline these potentials. We fixed the membrane potential at −60 mV and then we depolarized the cell at 0 mV during 10, 200, and 800 ms for 40 min to measure the amplitude of _f_I_AHP_, _m_I_AHP_, and _s_I_AHP_, respectively, at the end of each depolarized pulse in control conditions (ACSF) and after IGF-1 addition to the bath. The number of action potentials (spike frequency) elicited in response to a series of long (1 s) depolarizing current steps (25– 200 pA for basal condition and 25-350 pA for neurons recorded after the behavioral tests; 25 pA increments) were used to assess excitability of IL-L5PNs.

Bipolar stimulation was applied through a Pt/Ir concentric electrode (OP: 200 μm, IP: 50 μm; FHC) connected by 2 silver-chloride wires to a stimulator and a stimulus isolation unit (ISU-165 Cibertec). The stimulating electrode was placed at 100 μm below the soma of the recorded neuron (at the level of layer 6), close to the basal dendrites of the recorded IL-L5PNs. Paired pulses (100 μs in duration and 20–100 μA and 50 ms interval) were continuously delivered at 0.33 Hz. Excitatory postsynaptic currents (EPSCs) were isolated in the presence of GABA_A_R (50 μM picrotoxin; PiTX) and GABA_B_R (5 μM CGP-55845) antagonists. Inhibitory postsynaptic currents (IPSCs) were isolated adding AMPAR (20 μM CNQX) and NMDAR (50 μM D-AP5) antagonists. In both cases, after 5 minutes of stable baseline, we superfused IGF-1 for 35 min to check for long-term synaptic plasticity by analyzing the EPSCs and IPSCs peak amplitudes. In current-clamp mode, after a stable baseline of postsynaptic potentials (PSPs), the stimulation intensity was increased until suprathreshold responses reached were ≈21% for 5 min. Next, we applied IGF-1 for 15 min and measured the number of APs; afterward, we returned to initial stimulation intensity and measured the amplitude of PSPs to study the effect of IGF-1 on synaptic transmission. Miniature EPSCs (mEPSCs) were recorded at −60 mV in ACSF in the presence of 1 μM TTX, 20 μM PiTX, and 5 μM CGP-55845 to isolate excitatory synaptic transmission.

### Conditioned fear

Rats were anesthetized with ketamine (70 mg/kg i.p.; Ketolar™), xylazine (5 mg/kg i.p.; Rompum™), and atropine (0,05 mg/kg i.p.; B. Braun Medical S.A) and maintained with isoflurane (2-3% in oxygen). Animals were positioned in a stereotaxic apparatus (David Kopf Instruments, Tujunga, CA, USA) and placed on a water-heated pad at 37°C. The midline of the scalp was sectioned and retracted, and small holes were drilled in the skull. Rats were implanted with a single 26 gauge stainless-steel guide cannula (Plastics One) in the mPFC. Stereotaxic coordinates aiming toward the IL were (AP: 2.8 mm, LM: 1.0 mm, and DV: 4.1 mm) from bregma according to rat brain atlas (Paxinos and Watson, 2007), with the cannula angled 11° toward the midline in the coronal plane as described previously (Santini et al., 2004). A 33 gauge dummy cannula was inserted into the guide cannula to prevent clogging. Guide cannulas were cemented to the skull with dental acrylic (Grip Cement) and the incision was sutured. Buprenorphine hydrochloride (75 mg/kg s.c.; Buprex™) was administered for post-surgical analgesia. Rats were allowed at least 7 days for surgery recovery. Trace fear conditioning was conducted in a chamber of 25×31×25 cm with aluminum and Plexiglas walls (Coulbourn, Allentown, PA). The floor consisted of stainless-steel bars (26 parallel steel rods 5 mm diameter, 6 mm spacing) that can be electrified to deliver a mild shock. A speaker was mounted on the outside wall, and illumination was provided by a single overhead light (miniature incandescent white lamp 28 V). The rectangular chamber was situated inside a sound-attenuating box (Med Associates, Burlington, VT) with a ventilating fan, which produced an ambient noise level of 58 dB. The CS was a 4 kHz tone with a duration of 30 s and an intensity of 80 dB. The US was a 0.4 mA scrambled footshock, 0.5 s in duration, which co-terminated with the tone during the conditioning phase. Between sessions, floor trays and shock bars were cleaned with soapy water and the chamber walls were wiped with a damp cloth. An additional Plexiglas chamber served as a novel context for the auditory cue test. This chamber, which was a triangle with a black smooth Plexiglas floor, was physically distinct from the fear conditioning chamber. All the environmental conditions were completely different regarding the fear conditioning test (the intensity of light was lower, 10 V), except for the ventilating fan. Before placing rats into this chamber, chamber floor and walls were wiped with 30% vanilla solution to provide a background odor distinctive from that used during fear conditioning. The activity of each rat was recorded with a digital video camera mounted on top of each behavioral chambers. 30 min before extinction training, saline (NaCl 0.9%), IGF-1 (10 μM; Preprotech), or the IGF-1R antagonist 7-[*cis*-3-(1-azetidinylmethyl)cyclobutyl]-5-[3-(phenylmethoxy)phenyl]-7H-pyrrolo[2,3-]pyrimidin-4-amine (NVP-AEW 541, 40 μM; Cayman Chemicals) plus IGF-1 were infused into the IL. For the infusions, cannula dummies were removed from guide cannulas and replaced with 33 gauge injectors, which were connected by polyethylene tubing (PE-20; Small Parts) to 100 μl syringes mounted in an infusion pump (Harvard Apparatus). Drugs were infused at a rate of 0.5 μl/min for 1 min as described previously (Fontanez-Nuin et al., 2011).

#### Non-cannulated animals

Animals were divided into two groups, the conditioned group (COND) and the extinguished group (EXT). On day 1, rats received three habituation trials (tone-no shock; habituation phase) into two different contexts (context 1: square shock box and context 2: triangle box with soft floor). On day 2, rats received three conditioning trials (tone paired with shock; context 1; condition phase). On day 3, rats of COND group remained in their home cage, whereas rats of EXT group received twenty tone-alone trials (context 2; extinction phase). On day 4, both groups of rats received five tone-alone trials to test for recall of conditioning or extinction (test phase) (Figure supplement **3A**).

#### Cannulated animals

Animals were divided into the saline group (SAL) and the different drug groups (IGF-1 or NVP+IGF-1). On day 1, rats received three habituation trials into the two contexts aforementioned. On day 2, rats received three conditioning trials. On day 3, rats were infused with saline or drug 30 min before the beginning of the extinction phase. On day 4, all groups of rats received five tone-alone trials in the same chamber (context 2) to test for recall of extinction (Figure supplementupplement 3A). In all phases of the experiment, the interval between successive tones was variable, with an average of 2 min. All groups were tested on the same day to determine the long-term changes occurring in the mPFC that gate subsequent memory retrieval.

### Data analysis

Data were analyzed using pClamp (Molecular Devices), Excel (Microsoft), and GraphPad Prism 8.3 software. Twenty responses were averaged except when otherwise indicated. The magnitude of the change in peak EPSCs, IPSCs, and PSPs amplitude was expressed as a percentage (%) of the baseline control amplitude and plotted as a function of time. The presynaptic or postsynaptic origin of the observed regulation of EPSCs and IPSCs amplitudes was tested by estimating the paired-pulse ratio (PPR) changes, which were considered to be of presynaptic origin (Clark et al., 1994; Creager et al., 1980; Kuhnt and Voronin, 1994) and were quantified by calculating a PPR index (R2/R1), where R1 and R2 were the peak amplitudes of the first and second synaptic currents, respectively. To estimate the modifications in the synaptic current variance, we first calculated the noise-free coefficient of variation (CVNF) for the synaptic responses before and 40 min after applying IGF-1 in the bath with the formula CVNF = ✓δXPSC^2^ - δnoise^2^)/m; δXPSC^2^ and δnoise^2^ (X= E excitatory or X= I inhibitory) are the variance of the peak EPSC or IPSC and baseline, respectively, and m is the mean EPSC or IPSC peak amplitude. The ratio of the CV measured before and 40 min after applying IGF-1 (CVr) was obtained for each neuron as CV after IGF-1 responses/CV control (Clements, 1990). Finally, we constructed plots comparing variation in M (m after IGF-1 responses over m at control conditions) against the changes in response variance of the EPSC or IPSC amplitude (1/CVr2) for each cell. In these plots, values were expected to follow the diagonal if the EPSC or IPSC depression had a presynaptic origin. Results are given as mean ± SEM (N = numbers of cells). There were no gender differences in our experiments. MiniAnalysis software (SynaptoSoft Inc) and pCLAMP software were used for the analysis of the frequency and amplitude of mEPSCs. See Supplementary Table 1 for a detailed description of the statistical analyses performed in these experiments.

The total freezing time during the 30 s tone was measured and converted to percentage of freezing. Freezing was defined as the cessation of all movements except respiration. The percentage of freezing time was used as a measure of conditioned fear (Blanchard DC, 1972). The behavioral data were analyzed by manual evaluation of the videos and/or using the image J free software with the pulgging developed for this purpose (Shoji et al., 2014). No differences were observed with both methods. Values are reported as the means ± SEM. See Supplementary Table 1 for a detailed description of the statistical analyses performed in these experiments.

Blind experiments were not performed in the study but the same criteria were applied to all allocated groups for comparisons. Randomization was not employed. The sample size in whole-cell recording experiments was based on the values previously found sufficient to detect significant changes in past studies from the lab.

## RESULTS

### IGF-1 increases the excitability of IL-L5PNs

Ca^2+^-activated K^+^ currents that mediate the post-spike medium and slow afterhyperpolarization (mAHP and sAHP, respectively) are crucial in the regulation of neuronal excitability (Alger and Nicoll, 1980; Madison and Nicoll, 1984). We first tested the effect of IGF-1 on different components of afterhyperpolarizing potentials of IL L5PNs. In the current-clamp mode, we recorded mAHPs or sAHPs before and during bath application of IGF-1 (10 nM). IGF-1 reduced mAHP (Figure **1A,B**) and sAHPs (Figure **1C,D**), whereas the fast AHP (fAHP) was unaffected (Figure supplement **1A-B**). We also checked the effect on neuronal excitability by analyzing the number of actions potentials evoked by increasing current injection steps. We observed that the number of action potentials evoked by these protocols was higher during IGF-1 perfusion, indicating an increase in neuronal excitability mediated by IGF-1 (Figure **1E, F**).

**Figure 1.**
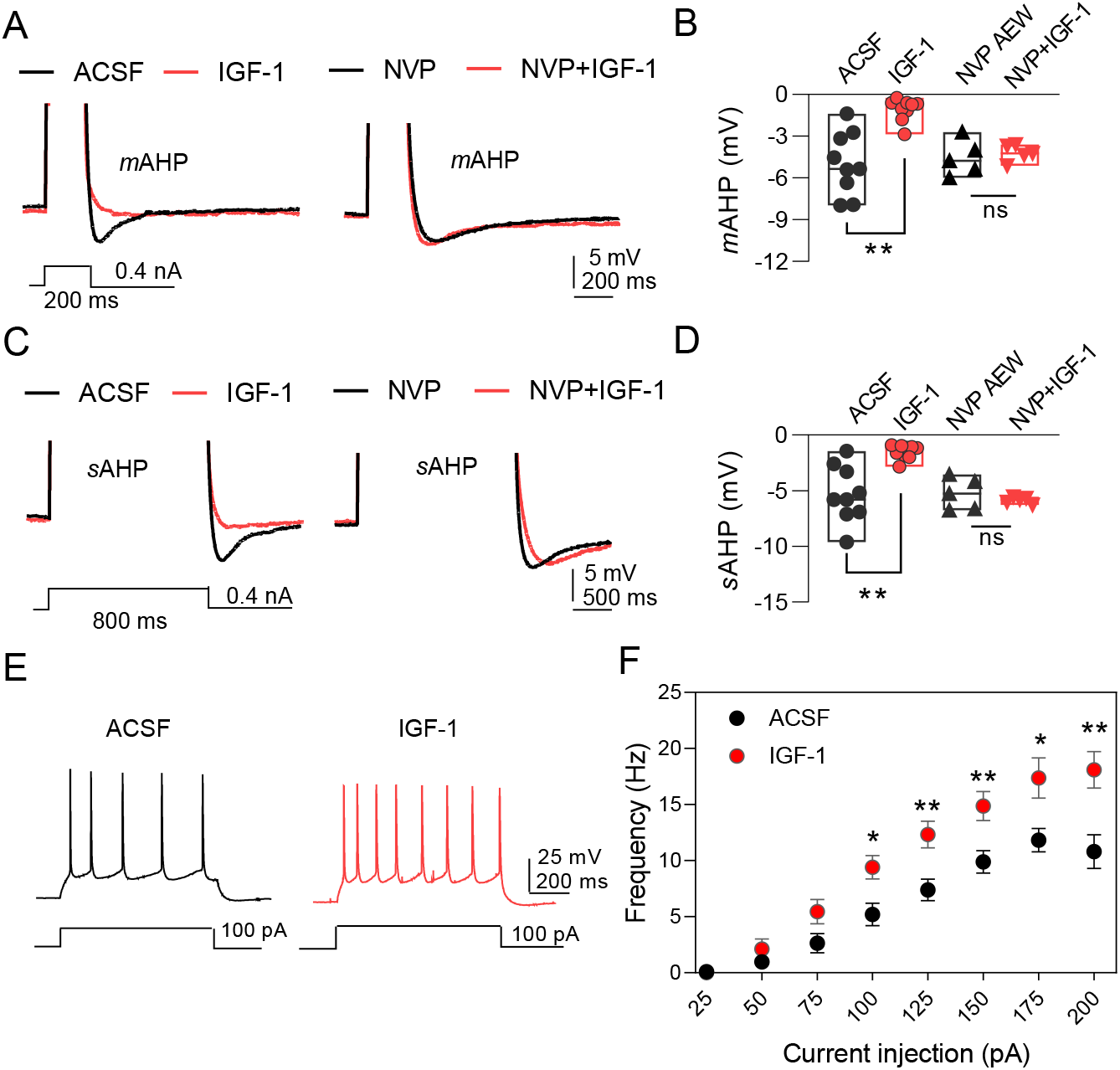
IGF-1 increases the excitability of IL-L5PNs. **A.** Representative recordings from IL-L5PNs of hyperpolarizing potentials elicited by a 200 ms depolarizing pulse to study medium AHP, in control conditions (ACSF, left) and in the presence of NVP-AEW541 (40 nM, right), before (black) and after of IGF-1 (10 nM, red) (spikes are truncated). **B.** Bar diagram summarizing mAHP amplitudes (n=5-9) ** p< 0.01, ns (non-significant) Student paired t-test. **C.** Same as A., but applying an 800 ms depolarizing pulse to study slow AHP. **D.** Bar diagram summarizing sAHP amplitudes (n=5-9) ** p< 0.01, ns (non-significant) Student paired t-test. **E**. Representative traces recorded from IL-L5PNs after 100pA current injection in ACSF (gray) and after IGF-1 application (red). **F.** Frequency-injected current relationships for IL-L5PNs in ACSF (gray) and after IGF-1 application (red) (n=8) * p < 0.05, Multiple t-test (Holm-Sildak methods). See also Figure 1—Figure supplement 1.

We next performed voltage-clamp recordings to analyze the effect of IGF-1 on the currents that underlie the mAHP and sAHP (Alger and Nicoll, 1980; Storm, 1990) (_m_I_AHP_ and _s_I_AHP_, respectively). IGF-1 also decreased _m_I_AHP_ (Figure **2A,B**) and _s_I_AHP_ (Figure **2D,E**), while the fast afterhyperpolarization current remained unaltered (Figure supplement **1C-D**). We observed a gradual current reduction that reached a plateau 20 min after IGF-1 application in both the _m_I_AHP_ and _s_I_AHP_ (Figure **2 C and F**). However, in the absence of IGF-1, the neuronal depolarization did not modify the AHP currents (Figure supplement **2A**). Interestingly, we observed a long-lasting reduction of sIAHP that remained after the wash-out of IGF-1 (Figure supplement **2B**) and required the activation of the AHPs by the depolarizing protocol (Figure supplement **2C**). Moreover, the IGF-1-dependent modulation of the _m_I_AHP_ and _s_I_AHP_ was prevented by the IGF-1R antagonist NVP-AEW541 (400 nM, Figure **2A-F**), indicating that these effects were mediated by the activation of the IGF-1R. Taken together, these results demonstrate that IGF-1 induced a long-term decrease of _s_I_AHP_ leading to an increase of neuronal firing frequency.

**Figure 2.**
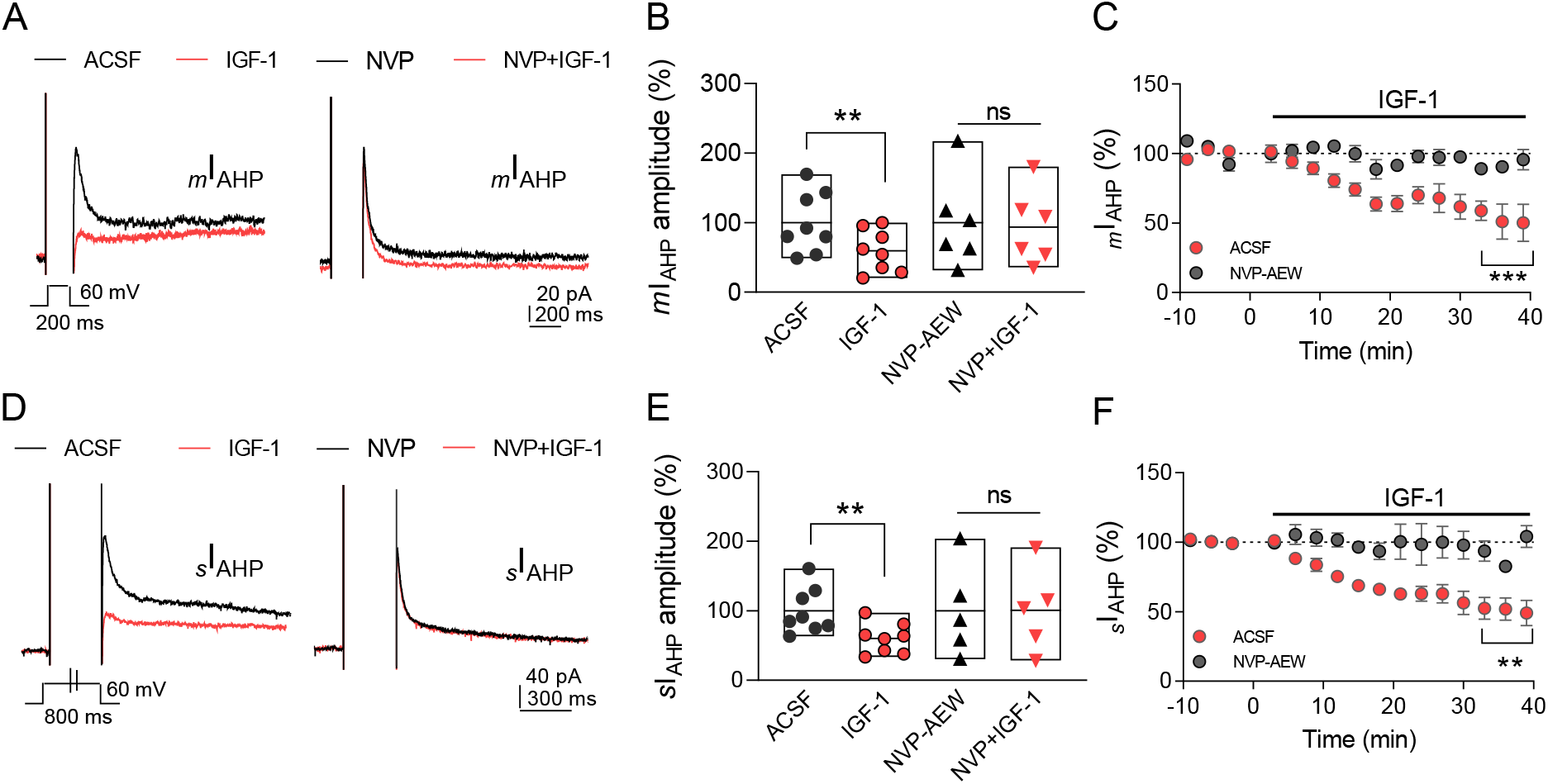
IGF-1 reduces _m_I_AHP_ and _s_I_AHP_ in IL neurons. **A.** Representative current traces recorded from IL-L5PNs in response to a 200ms depolarizing pulse from −60 mV to 0 mV in control condition (ACSF, left) and in the presence of (400 nM) NVP-AEW541 (right), before (black) and after addition of 10 nM IGF-1 (red). **B.** Normalized current amplitude of _*m*_I_AHP_ (n=5-8). ** p< 0.01, ns (non-significant) paired t-test. **C.** Time curse of _*m*_I_AHP_ in ACSF (red) and in presence of NVP-AEW541 (n=6-4) ***p<0.001 Student paired t-test. **D.** Same as A, but a 800ms depolarizing pulse was evoked. **E.** Same as B, but for _*s*_I_AHP_ (n=5-8) ** p< 0.01, ns (non-significant) paried t-test. **F.** Same as C, but for _*s*_I_AHP_ (n=6-5). See also Figure 2—Figure supplement 2.

### IGF-1 induces long-term potentiation of the synaptic transmission of IL-L5PNs

We next examined whether IGF-1 modulates the EPSCs. After isolating the EPSCs (see materials and methods), we measured their amplitude and analyzed the paired-pulse ratio and coefficient of variance. We found that IGF-1 induced long-term depression of EPSC peak amplitude (72.63% of baseline, Figure **3A**) that was prevented in the presence of NVP-AEW541 (97.72% of baseline, Figure **3A**). The IGF-1-mediated depression of the EPSCs was associated with changes in the paired-pulse ratio (PPR) (Figure **3B**) and coefficient of variation (1/CV^2^) (Figure **3C**), indicating a presynaptic mechanism. Additionally, this modulation was not observed when the synaptic stimulation was absent (Figure **3D**), suggesting that the evoked synaptic responses were required for this LTD of the EPSCs.

**Figure 3.**
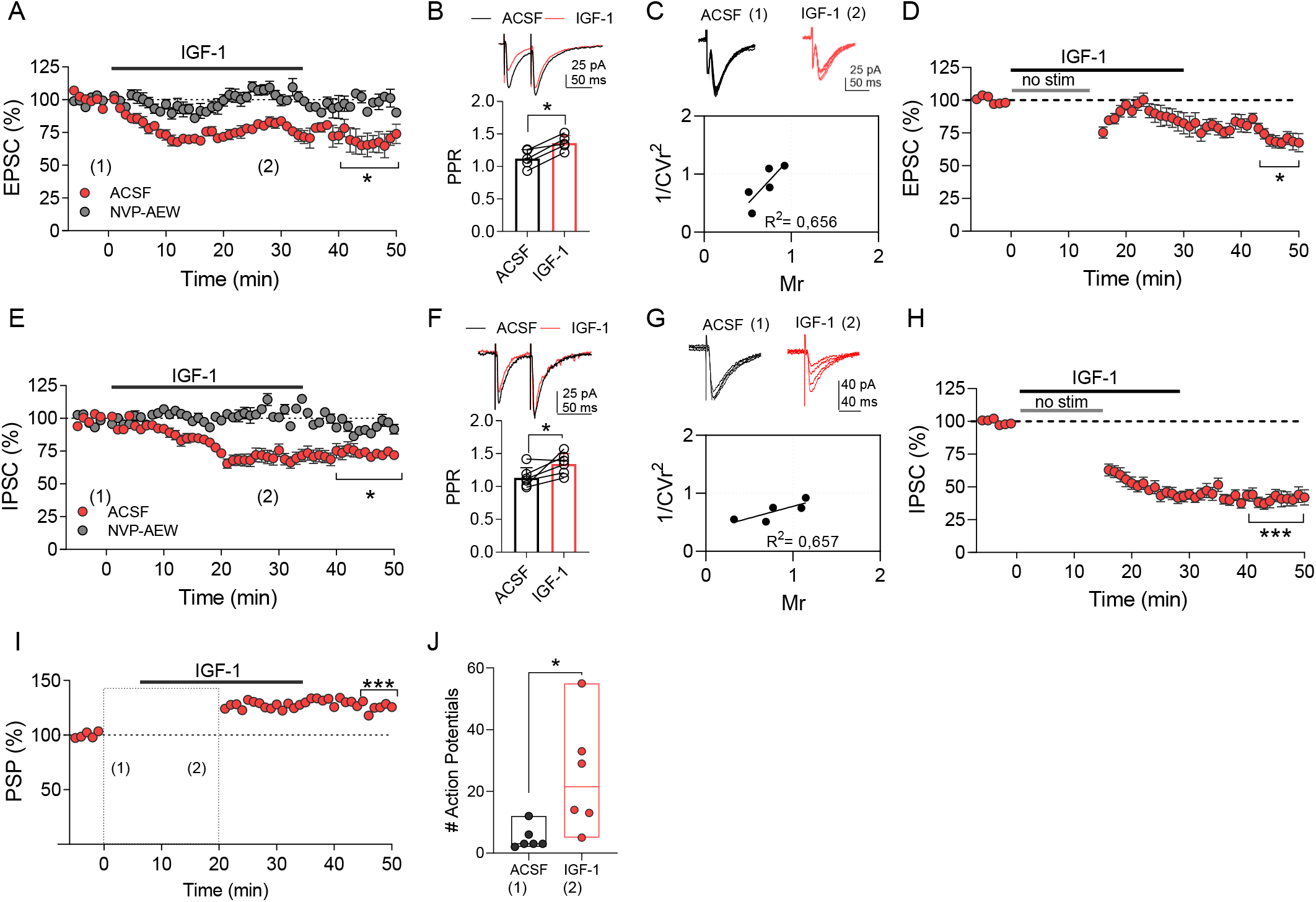
IGF-1 induces long-term potentiation in IL-PNL5. **A and E.** Time curse of EPSCs (n=5) or IPSCs (n=5) were recorded from IL-L5PNs in whole-cell voltageclamp mode. After a stable (~5-min) baseline, the IGF-1 was bath applied at zero time for 35 min. Student paired t-test *p>0.05. **B and F.** Representative traces, which correspond to the numbers in the time-course plot, are shown on the top. Bar diagrams summarizing the paired-pulse ratio (PPR) before and after IGF-1, on the bottom. Student paired t-test *p>0.05; (n=5). **C and G.** Representative traces are showed before and after IGF-1 on the top. Plot of the variance (1/CVr2) as a function of the mean EPSC or IPSC peak amplitude (M) 30 min after IGF-1 and normalized to control condition (ACSF) (n=5). **D and H.** Time curse of EPSCs or IPSCs showing long-term depression effect of IGF-1 in the absence of synaptic stimulus for 15 min. Student paired t-test *p>0.05 ***p>0.001 (n=6). **I.** Time curse of PSP were recorded from IL-L5PNs in whole-cell current-clamp mode. After a stable (~5-min) baseline, the IGF-1 was bath applied at zero time for 35 min. Student paired t-test ***p>0.001 (n=6). **J.** Bar diagrams summarizing the number of action potential in ACSF and after IGF-1 bath applied * p< 0.05 vs basal levels Student paired t-test (n=6).

We also analyzed whether IGF-1 regulates the IPSCs. We isolated the IPSC (see materials and methods) and after 35 minutes of IGF-1, we observed a long-term depression of IPSCs (41.99% of baseline, Figure **3E**) that were dependent of IGF-1R activation, since this effect was prevented by NVP-AEW541 (99.0% of baseline, Figure **3E**). The IGF-1-mediated depression of the IPSCs was associated with changes in the paired-pulse ratio (PPR) (Figure **3F**) and coefficient of variation (1/CV^2^) (Figure **3G**), pointing to a presynaptic mechanism. Nevertheless, the LTD of the IPSC did not require the evoked IPSCs since IGF-1 was able to induce it when the synaptic stimulation was absent (Figure **3H**).

Finally, we studied the effect of IGF-1 on the postsynaptic potentials (PSPs). After recording a stable baseline of PSPs, we increased the intensity of stimulus until that ≈21% of the responses recorded during 5 min were suprathreshold and then we added IGF-1 (Figure **3I** and **J**). IGF-1 induced a significant increase in the number of AP during 5 min of recording (Figure **3J**). When we returned to the basal intensity of stimulation, we observed a long-term potentiation (LTP) of the PSPs (Figure **3I**). Together, our results reveal that IGF-1 induces a presynaptic depression of both EPSCs and IPSCs, which triggers a LTP of PSPs.

### IGF-1 facilitates fear extinction

Since a reduction of the sAHP was previously shown to favor recall of extinction (Santini et al., 2008), we next analyzed whether IGF-1 was able to facilitate it. First, we performed a set of experiments to determine the number of sessions required to induced fear conditioning and extinction as previously described (Fontanez-Nuin et al., 2011) (Figure supplement 3). On day 2, the different groups of rats acquired similar levels of conditioned freezing (COND, 74%; EXT, 77%) with three sessions of association (CS-US, fear conditioning). On day 3, rats from the EXT group showed gradual within-session extinction across twenty trials to a final freezing level of 12% (CS-No US), while rats from the COND group remained in their home cage. On day 4, rats from COND group showed high levels of freezing (77-95%) on the test tones, whereas rats from EXT group showed low levels of freezing (50-35%), indicating good recall of extinction during five sessions (Figure supplement **3A-B**).

To study the effect of IGF-1 on fear extinction memory, we infused saline or IGF-1 or NVP+IGF-1 through a cannula guide previously implanted into the IL (Figure **4A**) 30 min before the extinction training (Figure **4B**). Although the level of freezing reached on day 2 was comparable in the three groups (Saline, 89%; IGF-1, 79%; NVP+IGF-1 81%), the rats from IGF-1 group showed less freezing in the last session on day 3 (Saline, 51%; IGF-1, 14%; NVP+IGF-1 59%). On day 4, IGF-1-treated rats exhibited less freezing than the saline and NVP+IGF-1 rats (Figure 4**B**). This result demonstrates that IGF-1 induces the facilitation of extinction memory dependent on the IGF-1Rs activation.

**Figure 4.**
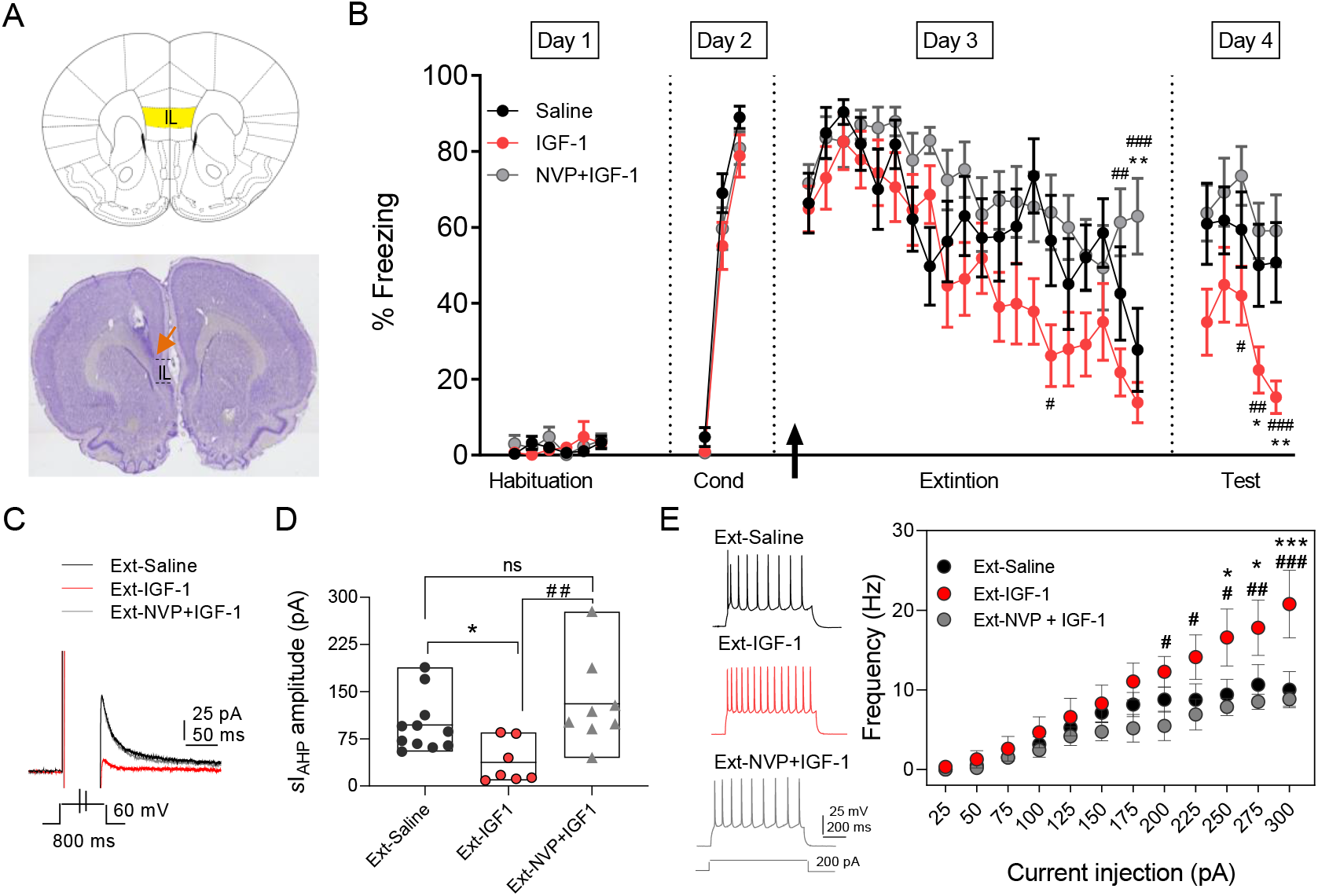
IGF-1 facilitates the memory of extinction by reducing the sI_AHP_ and increasing the firing frequency. **A.** (upper) Image adapted to atlas of brain rats showing infralimbic cortex (IL) localization. (bottom) Nissl-stained coronal section of rat infralimbic cortex, orange arrow indicates the tip of cannula implanted on the IL. **B.** Time curse of the percentage of freezing during the extinction protocol for the three groups studied. Saline (n=11), IGF-1 (n=14), and NVP-AEW541+IGF-1 (n=12) were directly applied into the IL through an implanted cannula 30 min before extinction training (day 3). The black arrow indicates the time of infusion. (*p<0.05 **p>0.01 Saline vs IGF-1 and NPV+IGF-1 #p>0.05; ##p>0.01; ###p>0.001 IGF-1 vs NPV+IGF-1; One-way ANOVA; Tukey’s multiple comparisons test). **C.** Representative sI_AHP_ current traces recorded from IL-L5PNs from animals that showed fear extinction **D.** Bar diagrams summarizing the _s_I_AHP_ for all the groups studied (n=6-10). Recordings were performed in day 4 (test) after finishing the extinction protocol; *p<0.05 Saline vs IGF-1 ##p>0.01 IGF-1 vs NPV+IGF-1 One-way ANOVA; Tukey’s multiple comparisons test. **E.** Representative traces recorded from IL-L5PNs from animals that showed fear extinction after 200 pA (left) current injection in Saline (black); IGF-1 (red) and NVP+IGF-1 (gray). Frequency-injected current relationships for IL-L5PNs in Saline (black) IGF-1 (red) and NVP+IGF-1 (gray). (*) IGF-1 vs. Saline and (#) IGF-1 vs. NVP+IGF-1; n=6-10; One-way ANOVA; Tukey’s multiple comparisons test). See also See also Figure 4—Figure supplement 3.

### IGF-1 induces an increase in neuronal excitability and a reduction in mEPSCs in rats exposed to the extinction memory task

Fear extinction is paralleled by an increase in the excitability of pyramidal neurons from layers 2/3 and 5 from IL (Santini et al., 2008). Therefore, we tested whether a change in the excitability of IL-L5PNs occurs in the animals in which IGF-1 facilitated the extinction memory. For this purpose, we recorded IL-L5PNs from all behavioral groups after the last extinction session on day 4. We observed that the _*s*_I_AHP_ amplitude of the IGF-1 group was smaller than the _*s*_I_AHP_ amplitude of the saline group (Figure **4C-D**). Importantly, this _*s*_I_AHP_ amplitude difference was not present when comparing the saline group with the group treated with NVP+IGF-1 (Figure **4C-D**). Moreover, the firing frequency of IL-L5PNs from the IGF-1-treated group was significantly increased compared with saline or NVP+IGF-1 groups. These results support that IGF-1 induces the facilitation of extinction through the reduction of _*s*_I_AHP_ and the increase of IL-L5PNs firing frequency.

Since we have demonstrated that IGF-1 induces a presynaptic LTD of the EPSCs (see Figure 3), we tested whether the excitatory synaptic transmission was depressed by IGF-1 in animals in which IGF-1 facilitated the extinction. For this purpose, we recorded miniature EPSCs (mEPSCs) in IL-L5PNs from all behavioral groups after the last extinction session on day 4. We observed that the mEPSCs frequency of the IGF-1 group was lower than in saline and NVP+IGF-1 groups (Figure **5A-C**), with no changes in the amplitude among groups (Figure **5A,D,E**). Taken together, these results demonstrate that IGF-1 facilitates the establishment of fear extinction memory, generating synaptic and intrinsic plasticity at IL-L5PN.

**Figure 5.**
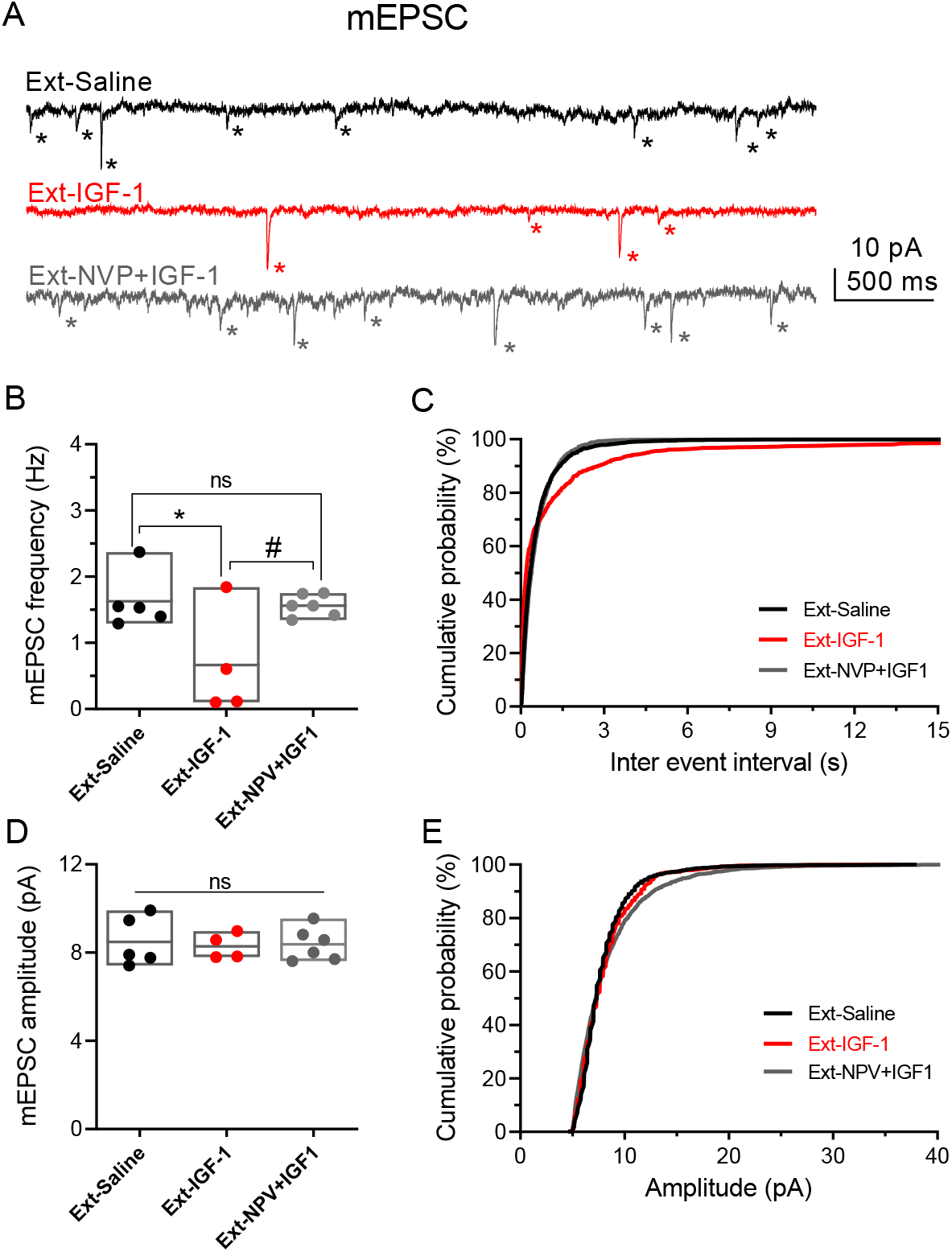
IGF-1 decreases the frequency of mEPSC. **A.** Representative currents traces recorded at −60 mV in IL-L5PNs from animals that showed fear extinction, in the presence of 1 μM TTX, 50 μM PiTX, and 5 μM CPG-55845 to isolate mEPSCs. Asterisks denote mEPSC events. Note the decreased mEPSCs frequency induced by IGF-1. **B.** Summary data (n=4-6) showing mean mEPSCs frequency from Ext-Saline (black); Ext-IGF1 (red) and Ext-NVP+IGF-1 (gray). One-way ANOVA; Tukey’s multiple comparisons test. **C.** Cumulative probability plots of mean inter-mEPSC interval in Ext-Saline (black); Ext-IGF1 (red) and Ext-NVP+IGF-1 (gray). Note that IGF-1 increased the mean inter-mEPSC interval (p < 0.05; Kolmogorov-Smirnov test). **D.** Summary data (n=4-6) showing mean mEPSCs amplitude. **E.** Cumulative probability plots of mean amplitude-mEPSC in Ext-Saline (black); Ext-IGF1 (red) and Ext-NVP+IGF-1 (gray). Note that IGF-1 does not change the mean mEPSCs amplitude (same cells as in C).

## Discussion

Interactions among the amygdala, the hippocampus, and the IL prefrontal cortex are important for the extinction of conditioned fear memory (Milad and Quirk, 2012; Orsini and Maren, 2012; Pape and Pare, 2010; Quirk and Mueller, 2008). Although the effects of IGF-1 on the amygdala (Stern et al., 2014) and the hippocampus (Chen et al., 2011; Stern et al., 2014) have been previously studied, there is no evidence about its possible actions on the IL. Our results demonstrate for the first time that IGF-1 applied to IL favors the extinction of conditioned fear memory causing an increase in L5PN excitability, by reducing the sAHP, and synaptic plasticity through the activation of the IGF-1R. We present new evidence showing that IGF-1 induced a long-lasting increase in excitability and LTP of the synaptic potentials at IL-L5PNs. The former is mediated by a significant reduction in the mAHP and sAHP, whereas the latter results from the interaction between presynaptic LTD of the EPSCs and IPSCs. Therefore, our results show a novel functional consequence of IGF-1 signaling on animal behavior in the mPFC. From this point of view, IGF-1 appears as a key endogenous molecule in the modulation of the extinction of conditioned fear memory, supporting the role of IGF-1 as a crucial piece in behavioral tasks.

There is a growing body of evidence supporting that fear extinction memory is encoded by IL neurons (Quirk and Beer, 2006) that show enhanced responses to extinguished cues during extinction recall (Milad and Quirk, 2002). Pharmacological (Hugues et al., 2004; Laurent and Westbrook, 2009; Sierra-Mercado et al., 2011; Sotres-Bayon et al., 2007), electrical (Milad et al., 2004), or optogenetic (Do-Monte et al., 2015) manipulations of IL have been described to modulate the acquisition of fear extinction. In addition, IL is involved in the recall of this memory since lesions of the IL produce deficits in its retention (Morgan and LeDoux, 1995; Quirk et al., 2000). In the present study, we infused IGF-1 into IL before the extinction protocol and found that the action of IGF-1 in these neurons induced a significant improvement of fear extinction memory compared to the groups treated with saline or NVP+IGF-1. Consistent with that fact, IGF-1 (LLorens-Martín et al., 2010; Trejo et al., 2008), and most recently also IGF-2 (Chen et al., 2011), have been related to cognitive function. Injections of IGF-1, IGF-2, or insulin into the amygdala did not affect memories critically engaging this region. However, bilateral injection of insulin into the dorsal hippocampus transiently enhances hippocampal-dependent memory whereas injection of IGF-1 has no effect (Stern et al., 2014). IGF-2 produces the most potent and persistent effect as a memory enhancer on hippocampal-dependent memories (Chen et al., 2011). Like insulin, IGF-2 did not affect amygdala-dependent memories when delivered into the BLA. Contextual fear extinction is facilitated when IGF-2 is injected into the dentate gyrus whereas inhibition of physiological IGF-1 signaling, via intrahippocampal injection of an IGF-1 blocking antibody, did not affect fear extinction (Chen et al., 2011). However, a single intravenous injection of IGF-1 before the training of contextual fear extinction increases the density of mature dendritic spines in the hippocampus and mPFC, favoring the memory of extinction (Burgdorf et al., 2017). All these reports are consistent with the idea that IGF-2 would have a specific role in the hippocampus while IGF-1 could have a selective role in the IL.

IGF-1 produces a strong potentiation of Ca^2+^ channel currents and the inhibition of A-type K^+^ channels in hippocampal and somatosensory neurons in the dorsal column nuclei, respectively (Blair and Marshall, 1997; Nuñez et al., 2003; Shan et al., 2003; Xing et al., 2006). Interestingly, our recordings from IL-L5PNs of animals treated with IGF-1 showed a long-term decrease of sAHP current, which is highly related to behavioral recall (Moyer et al., 1996; Santini et al., 2008), suggesting that IGF-1 also modulates the channels responsible for this current, at least in IL-L5PNs. Reduction in the amplitude and the duration of AHPs from IL-L5PNs caused by IGF-1 resulted in a significant increase in their excitability, as reflected in the increase of their firing frequency, a phenomenon that has been previously associated with fear extinction memory (Santini and Porter, 2010). Moreover, our results also suggest that IGF-1 could modulate the M-type K^+^ channels since the current generated by these channels mediate partially the _s_I_AHP_ and enhance the bursting of the IL during the acquisition of fear extinction (Santini and Porter, 2010). Since increases in excitability favors the induction of synaptic plasticity (Sepulveda-Orengo et al., 2013), our results suggest that IGF-1 would enhanced IL excitability, facilitating the induction of synaptic plasticity and improving the extinction memory (Santini and Porter, 2010).

Previous studies have demonstrated that the activation of metabotropic glutamate receptor type 5 (mGluR5) increase IL-L5PN excitability and favors the LTP of glutamatergic synaptic transmission, effects that are crucial in the recall of extinction (Fontanez-Nuin et al., 2011; Sepulveda-Orengo et al., 2013). Additionally, the administration of IGF-1 was reported to enhance synaptic plasticity in the hippocampus and the mPFC (Burgdorf et al., 2017; Ramsey et al., 2005). Therefore, we cannot rule out that an IGF-1-mediated increase in glutamatergic signaling could mediate both effects through the activation of mGluR5.

In conclusion, the present findings reveal novel mechanisms and functional consequences of IGF-1 signaling in IL. On the one hand, IGF-1 induces a reduction in sIAHP and increases the excitability of layer 5 pyramidal neurons of IL. On the other hand, IGF-1 modulates the excitatory and inhibitory synaptic transmission resulting in a long-lasting enhancement of the synaptic efficacy. Both the synaptic and intrinsic plasticity regulate the neuronal connectivity, which leads to the facilitation of consolidation of fear extinction memory. Altogether, these results strongly support the potential role of IGF-1 as a new therapeutic target for the treatment of anxiety and mood disorders.

## Acknowledgments

This work was supported by the following Grants:BFU2013-43668-P and BFU2016-80802-P AEI/FEDER, UE (MINECO) to D. Fernández de Sevilla.

## Author Contributions

Conceptualization, J.A.N-P., L.E.M., and D.FdeS.; Methodology, J.A.N-P., L.E.M., and D.FdeS.; Formal analysis, J.A.N-P., L.E.M., I.B.M., J.M-C., and A.M-C.; Investigation, J.A.N-P., L.E.M., I.B.M., J.M-C., A.M-C., and M.C.; Writing-Original Draft, L.E.M., and D.FdeS.; Writing-Review & Editing, J.A.N-P., L.E.M., I.B.M., and D.FdeS.; Funding Acquisition, D.FdeS.

---

**Figure supplement 1.**
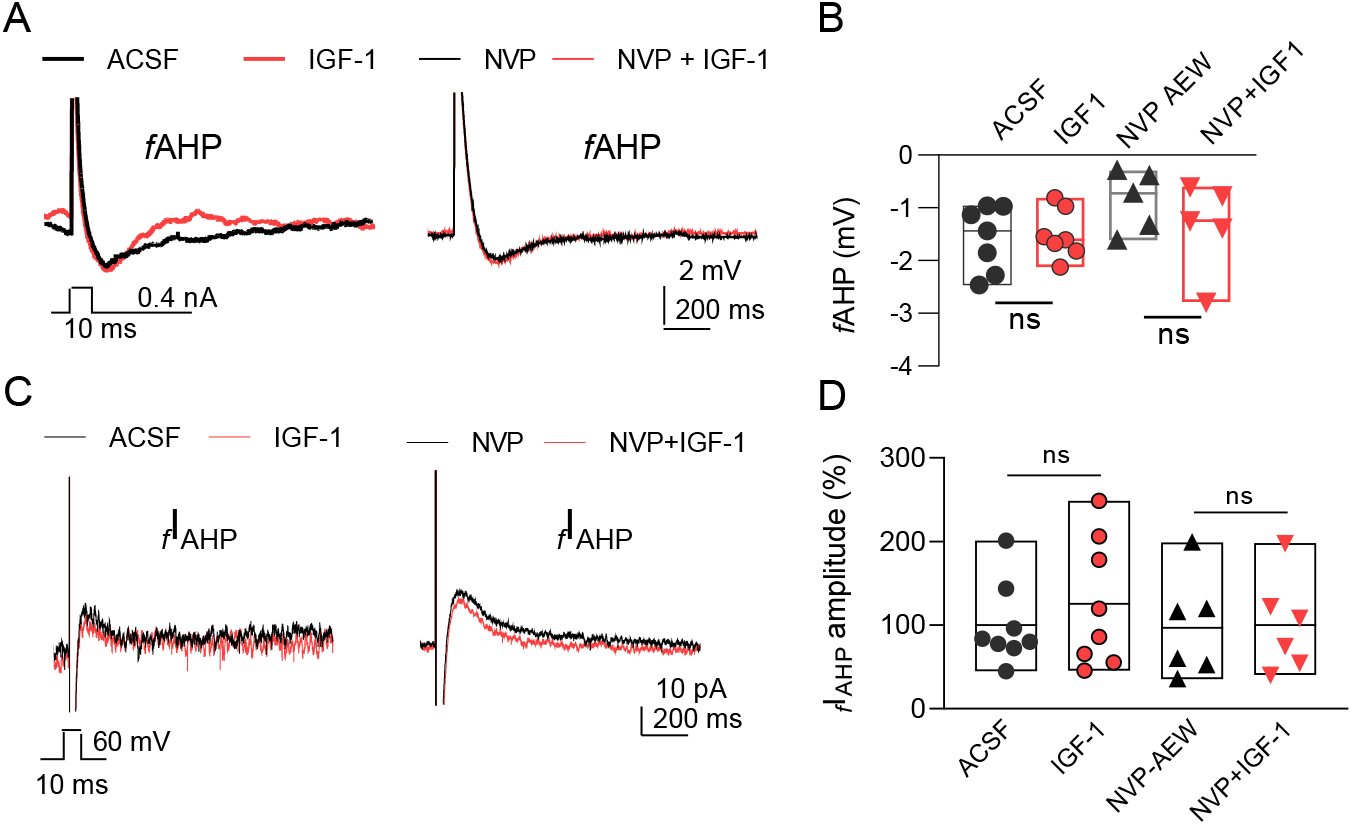
IGF-1 has no effect on *f*AHP and _f_I_AHP_. **A.** Representative recordings from IL-L5PNs of hyperpolarizing potentials elicited by a 10-ms depolarizing pulse to study the fast components of the AHP, in control conditions (ACSF, left) and in the presence of NVP-AEW541 (right), before (black) and after addition of IGF-1 (red) (spikes are truncated). **B.** Bar diagram summarizing *f*AHP (n=5-8) ns (non-significant), Student paired t-test. **C.** Representative current traces recorded from IL-L5PNs in response to a 10ms depolarizing pulse from −60 mV to 0 mV in control condition (ACSF, left) and in the presence of (400 nM) NVP-AEW541 (right), before (black) and after addition of 10 nM IGF-1 (red). **D.** Normalized current amplitude of _*f*_I_AHP_ (n=5-8) ns (non-significant) Student paired t-test.

**Figure supplement 2.**
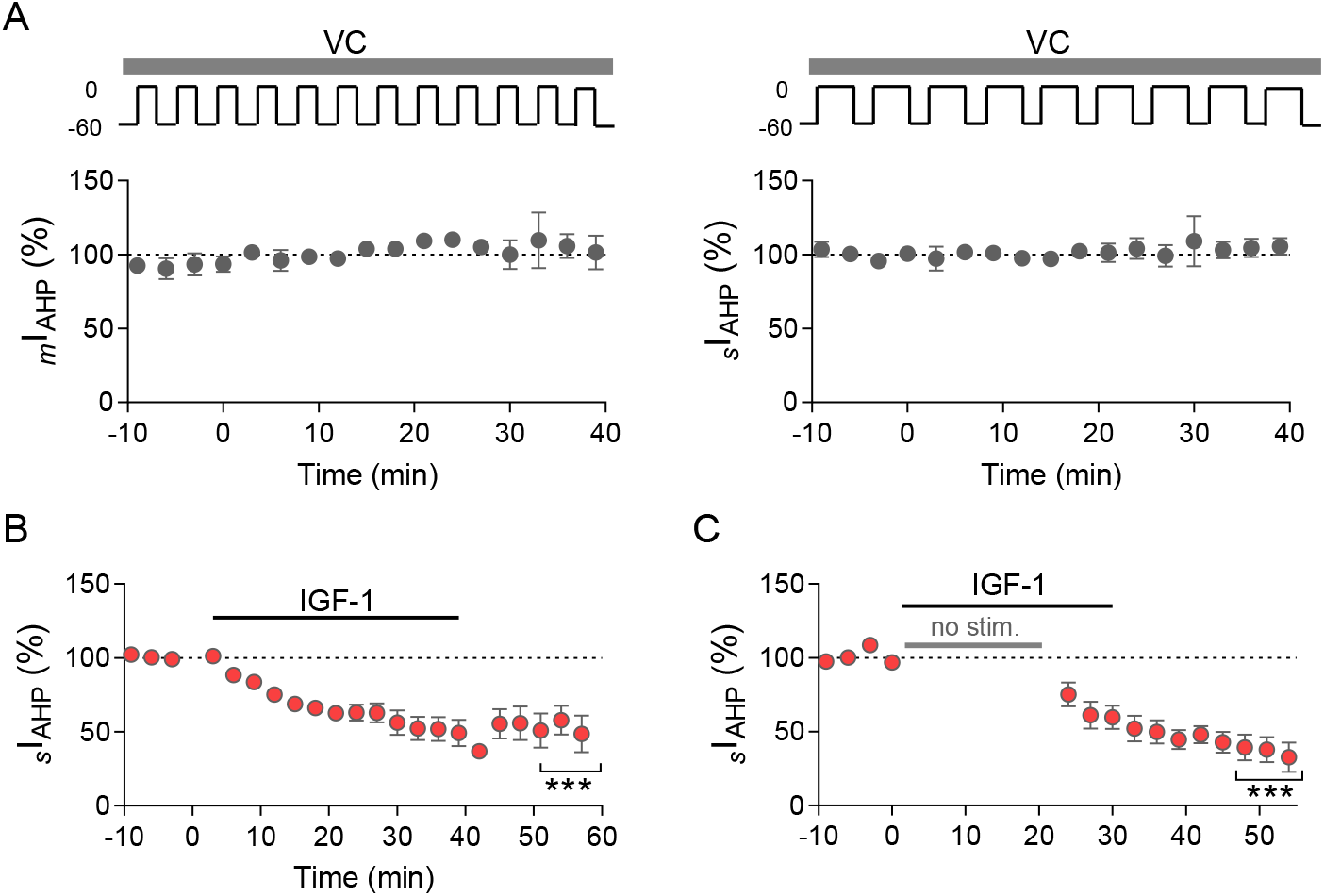
Time curse of _m_I_AHP_ and _s_I_AHP_. **A.** Time curse of _m_I_AHP_ (left) and _s_I_AHP_ (right) in control conditions (ACSF) after applying 200 and 800 ms depolarizing pulses, respectively (n=4 in each group). Note the protocol applied no change the currents **B.** Extended time curse of _s_I_AHP_ showing the persistent effect of IGF-1 after washout (n=8) *** p<0.001 vs basal levels; Student paired t-test. **C.** Time curse of _s_I_AHP_ showing long term depression effect of IGF-1 in the absence of the 800-ms depolarizing protocol (n=6) ***p<0.001 vs basal levels; Student paired t-test.

**Figure supplement 3.**
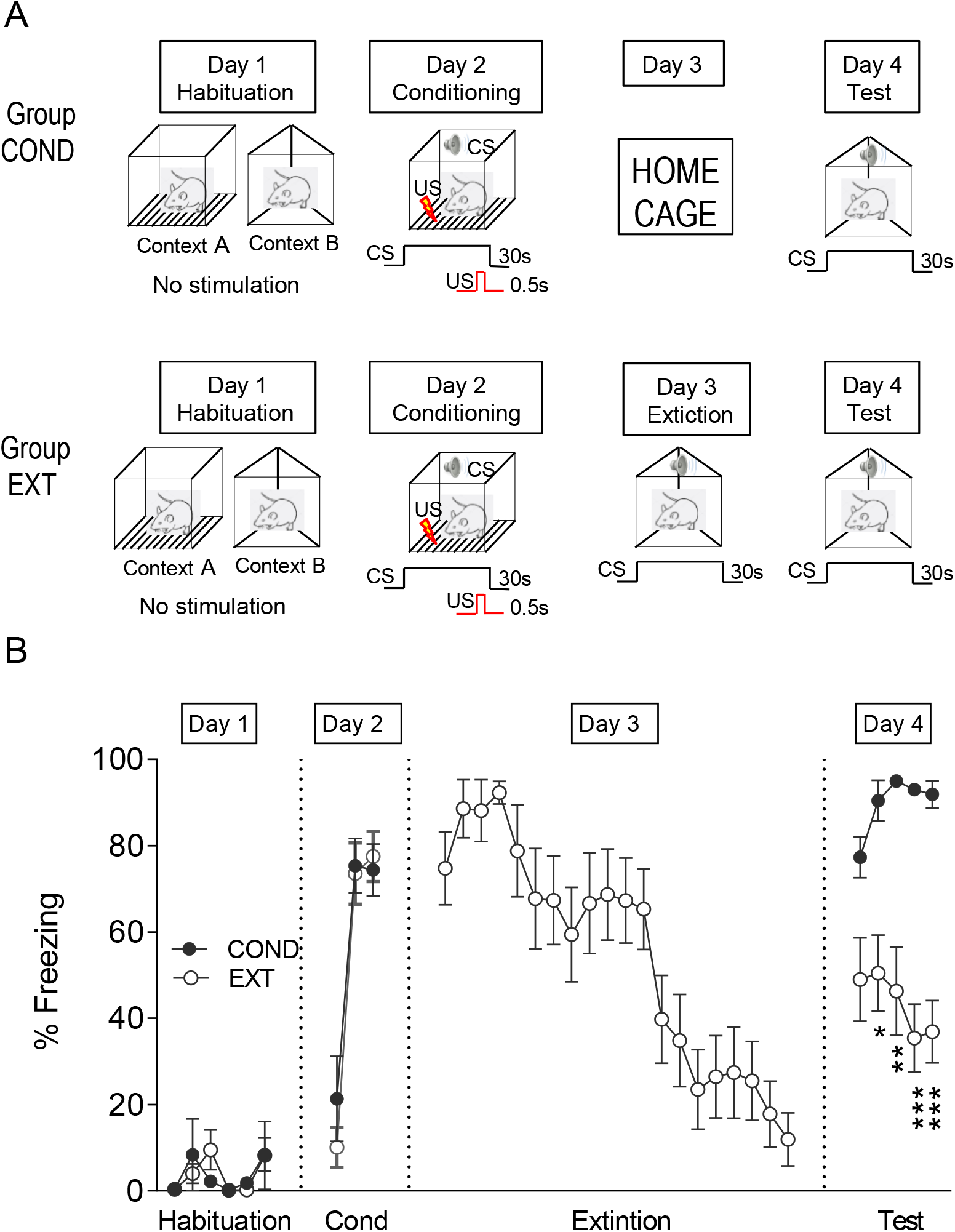
Protocol of behavior. **A.** Experimental design. **B.** Time curse of the percentage of freezing during the experiment for the two groups studied: conditioned (COND; n=7) and extinction (EXT; n=11) groups. As expected, animals in the COND group showed higher levels of freezing to the test tones compared with the EXT groups (***p<0.001; **p<0.01; One-way ANOVA, Tukey’s multiple comparisons test).

